# The persistent influence of caste on under-five mortality: Factors that explain the caste-based gap in high focus Indian states

**DOI:** 10.1101/516070

**Authors:** Jayanta Kumar Bora, Rajesh Raushan, Wolfgang Lutz

## Abstract

**Objective:** Although under-five mortality (U5M) is declining in India, it is still high in a few selected states and among the scheduled caste (SC) and scheduled tribe (ST) population of the country. This study re-examines the association between castes and U5M in high focus Indian states following the implementation of the country’s National Rural Health Mission (NRHM) program. In addition, we aim to quantify the contribution of socioeconomic determinants in explaining the gap in U5M between the SC/ST population and non-SC/ST population in high focus states in India.

**Data and method:** Using data from the National Family Health Survey (NFHS), we calculated the under-five mortality rate (U5MR) by applying a synthetic cohort probability approach. We applied a binary logistic regression model to examine the association of U5M with the selected covariates. Further, we used Fairlie’s decomposition technique to understand the relative contribution of socioeconomic variables on U5M risk between the caste groups.

**Findings:** In high focus Indian states, the parallel gap in U5M between well-off and deprived caste children has disappeared in the post-NRHM period, indicating a positive impact in terms of reducing caste-based inequalities in the high focus states. Despite the reduction in U5M, particularly among children belonging to STs, children belonging to the SC and ST population still experience higher mortality rates than children belonging to the non-SC/ST population from 1992 to 2016. Both macro level (district level mortality rates) and individual (regression analysis) analyses showed that children belonging to SCs experience the highest likelihood of dying before their fifth birthday. A decomposition analysis revealed that 78% of the caste-based gap in U5M is due to the effect of women’s level of educational attainment and household wealth between the SC/ST and non-SC/ST population. Program indicators such as place of delivery and number of antenatal care (ANC) visits also contributed significantly to widening caste-based gaps in U5M.

**Conclusion:** The study indicates that there is still scope to improve access to health facilities for mothers and children belonging to deprived caste groups in India. Continuous efforts to raise the level of maternal education and the economic status of people belonging to deprived caste groups should be pursued simultaneously.

## Introduction

Mortality among children of age five and below has declined in most countries, with the decline accelerating since mid-2000 [1,2]. Yet, in the era of the Sustainable Development Goals (SDGs), the issue continues to be a major public health concern, particularly in low- and middle-income countries (LMICs). Among the LMICs, India contributes the highest number of deaths in children under five and socially disadvantaged groups disproportionately carries the burden of children dying at a young age.

The burden of child mortality in high focus states of India has been the concern of policy makers and researchers equally. The high-focus states in India were designated as such by the Indian government because of their persistently high child mortality and relatively poor socio economic and other health indicators. A recent study demonstrated that the majority of the districts in the high focus states are not likely to achieve the SDGs concerning the preventable death of new-borns [3]. This study re-examines the association between castes and under-five mortality (U5M) in high focus Indian states using the most recent data. It also aims to quantify the relative contribution of socioeconomic determinants to U5M by explaining the gap between socially disadvantaged (scheduled castes and scheduled tribes) and non-disadvantaged castes in high focus states.

### Caste affiliation and under-five mortality in India

The Indian caste system is a traditional system of social stratification that has existed for more than three thousand years [4]. It is a social stratification system of self-governing and closed groups or communities called Jatis. These Jatis are assigned by birth and remains the same throughout an individual’s life. The ancient Varna system divided Hindu society into initially four, and later five, distinct Varna or castes, that are mutually exclusive, hereditary, endogamous and occupation-specific. These are the Brahmins (priests), Kshatriyas (warriors), Vaisyas (traders and merchants) and Sudras (those engaged in menial jobs) and those doing the most despicable menial jobs-the Ati Sudras or the former untouchables [5]. The ‘untouchables’ have the lowest social standing. Another way of categorizing the castes, which is now used in India to direct certain policies, is scheduled castes (SCs), scheduled tribes (STs), other backward classes (all disadvantaged groups) and general castes (non-disadvantaged castes). People who belong to SCs were previously referred to as “untouchables”, while the STs are communities of people living in tribal areas (mainly forest). SCs and STs are historically marginalized and disadvantaged social groups and are officially recognized and listed by the Indian Constitution. Interestingly, although castes originated within the Hindu religion, it also exists in the other religious groups in India such as Islam and Christian [6]. According to data obtained from the 2011 census of India, together, they constitute 25.2% of the country’s total population (with SCs contributing 16.6% and STs 8.6%).

Despite continuing efforts by modern governments to redress the effects of the caste system through a system of reservation (positive discrimination), caste remains a significant line of social division in India. The Indian Constitution has given people belonging to disadvantaged groups a special status since 1950 and makes provision for quotas in politics, education and job opportunities, as well as various other arrangements, including laws to abolish practices prolonging social inequities and development programs specially designed to cater to the needs of these groups [7]. However, they continue to face multiple difficulties compared to the rest of the population [8–10] and still have lower socioeconomic development indicators than the rest of the population [11]. These people are generally exposed to poor living conditions, observe a poor diet and have limited access to health care. In addition to their low socioeconomic circumstances, people in disadvantaged castes experience other adverse circumstances such as caste-based discrimination while accessing the health care system in India [12]. In addition, their life expectancy is relatively low and both child and adult mortality are relatively high [13]. In fact, people belonging to disadvantaged castes constitute almost 50% of all maternal deaths in the country and their children are more undernourished compared to the rest of the population [14,15].

The association between caste and child mortality is well documented [13,16–24]. In general, previous studies showed that children belonging to disadvantaged castes such as SCs/STs experience a higher likelihood of death compared to children belonging to non-deprived castes. It has also been found that caste differences in infant and child mortality are substantially reduced when parental socioeconomic characteristics are held constant [21]. In a recent study by Ranjan et al. (2016) [25], the authors concluded that the gap in infant mortality between tribal and non-tribal populations was substantial in the early months after birth, narrowed between the fourth and eighth months, and grew thereafter. The study by Dommaraju and colleagues (2008) [17] examined the effect of caste on child mortality and maternal health care utilization in rural India. They concluded that children belonging to lower castes have a higher risk of death and that women belonging to the lower castes have lower rates of antenatal and delivery care utilization than children and women belonging to the upper castes. The study further suggested the need to target low-caste members in the provision of maternal and child health services.

Has the association between caste and under-five mortality however been fading away in recent years due to the government’s health programs? In 2005, the Indian government launched the National Rural Health Mission (NRHM), which was renamed the National Health Mission (NHM) in 2013, in an effort to improve the availability of and access to quality health care for the poor, as well as for women and children, especially in rural areas. The program was introduced with a special focus on the nine socioeconomically disadvantaged high focus states. In the same period, the reproductive, maternal, newborn, child and adolescent health (RMNCH+A) approach was launched to address major causes of mortality among women and children, as well as the delays in accessing and utilizing health care and services. Many components of the NHM directly address the issues related to U5M and health status. There have been numerous initiatives under the NRHM-NHM to enhance newborn care and delivery. Through the introduction of the Janani-Shishu Suraksha Karyakram (JSSK), the Accredited Social Health Activists (ASHA) are expecting to increase institutional delivery and significantly bring down the neonatal mortality rate. The Indian government is implementing Janani Suraksha Yojana (JSY) which is a safe motherhood scheme to reduce maternal and infant mortality by promoting institutional delivery among pregnant women by providing conditional cash assistance. The larger institutional framework of NHM complements the JSY cash incentive by providing comprehensive healthcare, including antenatal and post-natal services, transport to facilities, and support services from ASHA. It includes several support services administered by community health workers to encourage pregnant women to use healthcare facilities for childbirth, along with at least three antenatal check-ups [26].

Following the implementation of NRHM, India has avoided nearly one million child deaths across socioeconomic groups between 2005 and 2015 [27]. The program is credited with having greatly reduced the inequities in maternal health services through increased institutional delivery and antenatal care in the high focus and deprived states of India [28].

Does this substantial coverage of maternal and health care services in recent years however reduce the caste-based gaps in U5M in high focus states? Recently available nationally representative data, commonly known as the National Family Health Survey (NFHS) allowed us to re-examine the association between caste and child mortality in India. We also wanted to determine what the relative role of socioeconomic characteristics was in explaining the gap in U5M between the caste groups in the country, if the caste-based gap in U5M were to continue. To our knowledge, these questions had not previously been answered in recent literature. Therefore, we aimed to extend the previous knowledge on caste disparity in several directions. First, we re-examined the association between caste and U5M in the high focus states. Our study is exclusively based on high focus states in north-central and eastern India, which contributed nearly 46% of the total number of under five deaths between 2005 and 2015 and were found to be lagging behind in terms of achieving the SDG goals on U5M [3]. Secondly, we provide district level estimates of U5M for the SC and ST population. To our knowledge, no previous study has examined district level variations in the under-five mortality rate (U5MR) among disadvantaged castes. Finally, we explain the caste based gap in U5M by applying an extension Oaxaca-type decomposition for nonlinear models as suggested by Fairlie [29]. The findings of this study can help to understand the factors behind the pervasive gap in U5MR between deprived and other caste groups in India.

## Data

### Ethics statement

The study is based on an anonymous publicly available dataset with no identifiable information on the survey participants and no ethics statement is required for this research work.

We used data from the fourth round of the Indian Demographic Health Surveys (DHS), commonly known as National Family Health Surveys (NFHS) conducted between 2015 and 2016. Data from the previous three rounds of NFHS were only used for trend analysis. The 2015-2016 NFHS survey was conducted by the International Institute of Population Sciences (IIPS), Mumbai under the stewardship of the Ministry of Health and Family Welfare (MoHFW) of the government of India. The survey was based on 1,315,617 births from 601,509 households and was covered by 699,686 interviews with women aged between 15 and 49 years old. The sample was selected using a two-stage sample design and covers all 640 districts as per the 2011 census. During the first stage, villages was selected as the primary sampling units (PSUs) for rural areas with probability proportional to size, while for urban areas, census enumeration blocks (CEB) were used. During the second stage, a random selection of 22 households in each PSU and CEB was made for rural and urban areas, respectively. The unit level data is available from the DHS data repository and can be accessed on request. A detailed description of the survey design of the NFHS-4 is available in the national report [30].

We restricted our analysis to the high focus states that used in previous studies [31–34] with women belonging to 304 districts according to 2011 census of India. The nine high focus states are Assam, Bihar, Chhattisgarh, Jharkhand, Madhya Pradesh, Odisha, Rajasthan, Uttarakhand and Uttar Pradesh. These are the most populous, containing 48.5% of India’s population. They are located in the north-central and eastern belt of the country, are characterized by poor demographic and health indicators, and have under five and neonatal mortality rates higher than the national level.

## Methods and measures

### Outcome variable

The outcome variable in the study is U5M, which is defined as the probability of dying before reaching the age of five. We assigned a value of “1” if the child died and “0” if the child was alive for the outcome variable.

### Predictors

The predictors used in this analysis are broadly divided into three categories: demographic, socioeconomic and program variables. These variables were considered as they were found to be important determinants of U5M in previous literature. Under demographic variables, we included the sex of the child, the mother’s age at the birth of her first child (<20, 20-24 and 25 years or more) and the birth order of the children (1, 2-3 and 4+).

Under socioeconomic variables, we included caste affiliation, parents’ level of educational attainment, type of residence and wealth quintile of the household, type of fuel used for cooking, type of toilet facilities and source of drinking water. The caste group is the core predictor used in this analysis. We categorized caste into three categories namely SC, ST and non-SCST. We categorized the mother and father’s level of education into three groups: primary or below (5 or less than 5 years of schooling), secondary (6 to 12 years of schooling) and higher (more than 12 years of education).

In terms of wealth, the NFHS survey did not collect information on income. The economic status of a woman was assessed by computing a composite index of household wealth indicating possession of wealth or assets by the household to which they belonged. We computed the wealth quintile of the household separately using the methodology followed in the fourth round of the NFHS. We excluded the variables of sanitation, source of drinking water and cooking fuel while constructing the wealth index. Using the total score, a household was categorized as belonging to the poorest, poorer, middle, richer and richest group. For convenience of analysis we grouped the five categories of the wealth quantile into three simplified categories by combining the poorest and poorer group into a new category named “poor“, the middle group remained “middle’ and the richer and richest groups were combined to form the “rich” category. Among the program indicators, the place of delivery of the child (home vs institutional delivery) and frequency of antenatal care (ANC) visits (none, 1 time, 2-3 times and 4+) during the pregnancy of last birth was considered in our analysis.

## Measures

### U5MR estimation

Mortality estimation was carried out using a dataset with full birth histories and information gathered from women aged 15 to 49 years surveyed in the NFHS. In the analysis, we considered only those births and deaths that took place five years preceding the survey. However, we chose the reference period of the district level U5MR estimates as “ten years prior to the survey date” in order to maximize the size of the sample needed to estimate those rates at district level. We applied direct methods of mortality-rate estimation using data on the children’s date-of-birth and their survival status, as well as the date-of-death and age at death of deceased children. Rustein and Rojas describe the synthetic cohort probability approach using the full birth history data of women aged 15 to 49 to estimate the U5MR rate [35] on DHS data. A synthetic cohort life table approach used in a previous study (3) used the same method as that used to produce the child mortality rate in DHS reports using the STATA package “syncmrates” [36].

We carried out the district level estimations of the U5MR for SCs, STs and non-SC/ST populations separately, with a 95% confidence interval and level of significance. Our inferences regarding the U5MR are strictly based on the districts for which 1) estimated U5MR are statistically significant at a 10% level of significance; and 2) estimated mortality rates are greater than zero. Thus, we provide all estimates in the appendices to this paper, and showed significant, insignificant, and “not enough sample size” in the districts for estimates separately in Figure 2.

**Fig 1.**
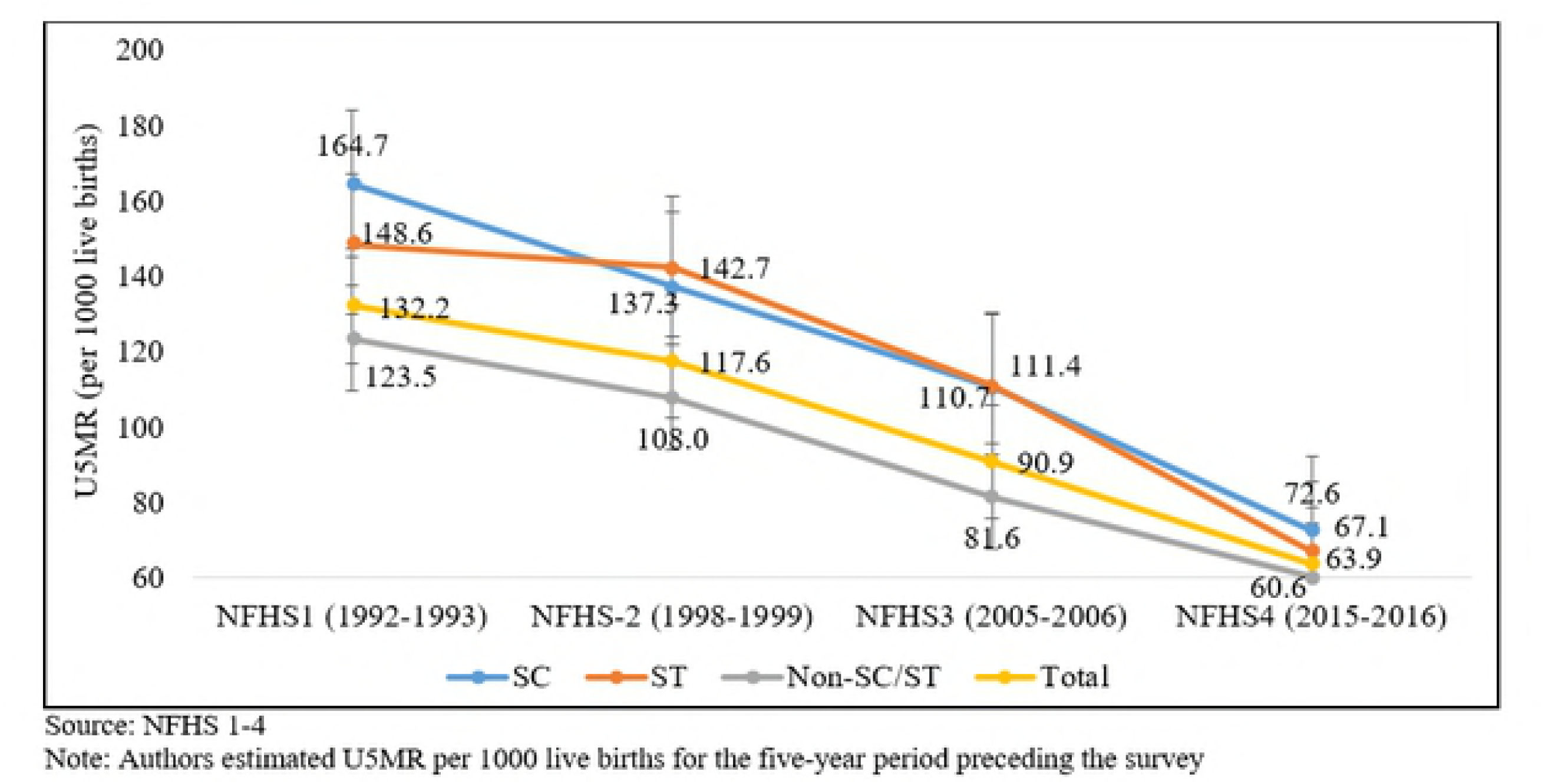
Trends in U5MR among social groups in high focus states, India 1992-2016

**Fig 2.**
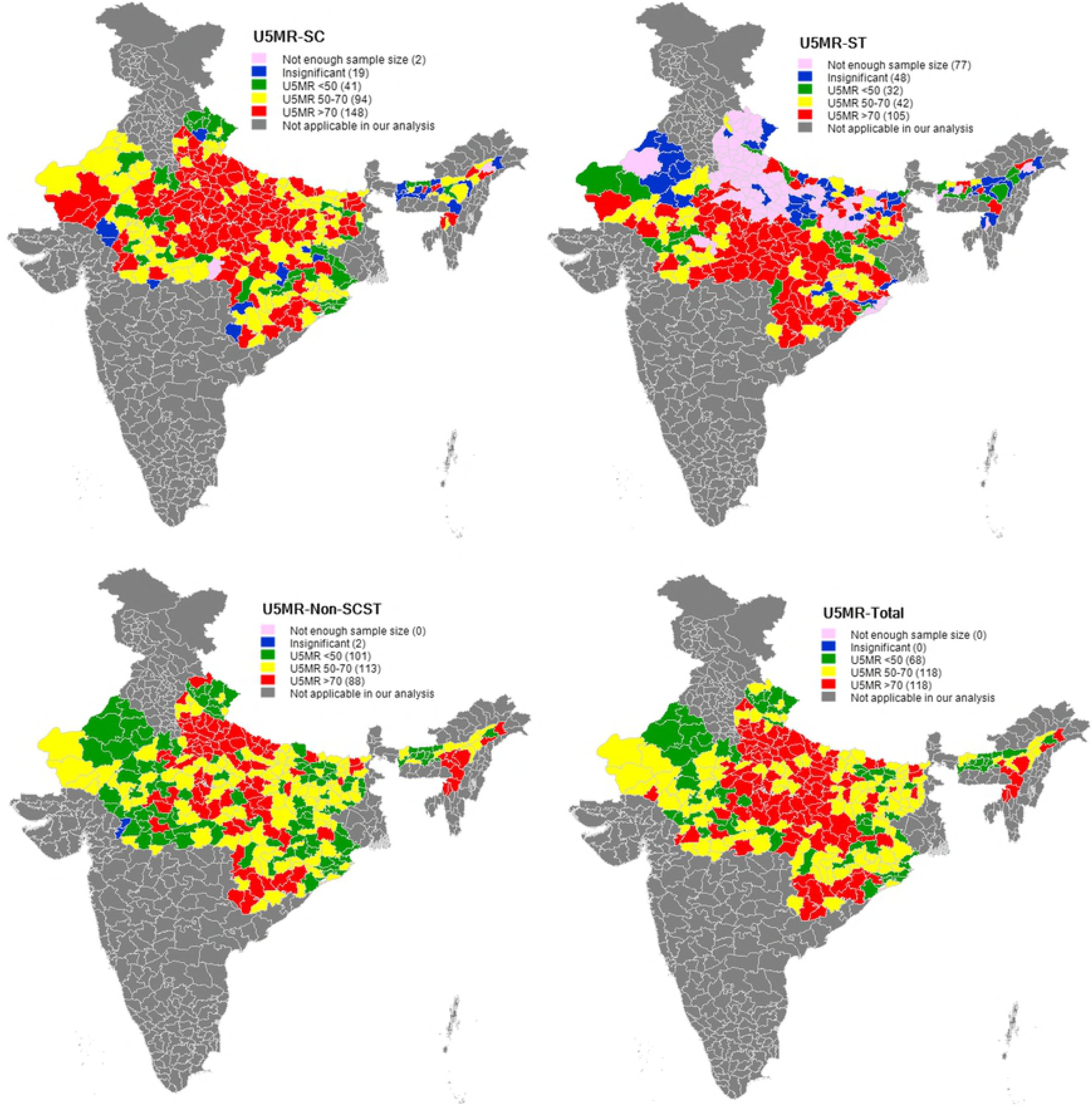
District wise U5MR (per 1000 live births) for SC, ST, non-SC/ST and Total populations in high focus states of India, 2015-16

## Statistical analysis

We used bivariate analysis to examine the differences in outcome and selected predictors between the SC, ST and non-SC/ST population. A binary logistic regression model was employed to examine the association between U5M and exposure variables. All exposure variables were tested with the variance inflation factor (VIF) to account for possible multi-collinearity before using the binary logistic regression model.

## Decomposition analysis

The Blinder–Oaxaca decomposition technique [37,38] is commonly used to identify and quantify the factors associated with inter-group differences in the mean level of outcome. This technique, however, is not appropriate if the outcome variable is binary, such as child mortality. Hence, we used the extension of the Blinder–Oaxaca technique developed by Fairlie [29] which is appropriate for binary models, to decompose the gap between social groups with U5M risk into contributions that can be attributed to different factors. For the decomposition analysis, we used the *‘fairlie’* command available for Stata. A detailed description of this method is discussed in appendix (Appendix S1). We used STATA S.E. 15.0 (STATA Corp., Inc., College Station, TX) version software to carry out the mentioned analysis in this study.

## Results

### Level and trends in U5MR among different social groups in high focus states of India

Fig 1 shows the trends in U5MR within the SC, ST and non-SC/ST populations in high focus states together for 1992-93 to 2015-16. It is observed that despite U5M having declined the most among children belonging to STs, it is still higher among SCs and STs than among the non-SC/ST population during 1992 to 2016. Compared to 1992-2005, the reduction in the U5MR was higher in the most recent period irrespective of caste groups. It is also clear from Figure 1 that the U5MR in high focus states is substantially higher than the target specified in SDG3 (at least as low as 25 per 1000 live births in 2030) and the national average (50 deaths per 1000 live births in NFHS4) for preventable deaths among children under five. On average, the estimated U5MR of high focus states are more than double the amount specified in the SDG3 target across all caste groups. Another striking point of this graph is that the parallel gap in U5MR between SC and ST and non-SC/ST populations between 1992 and 2005 was reduced drastically in 2015-2016.

Table 1 shows the current level of U5MR of SC, ST and non-SC/ST population in the selected high focus states. The estimation result is presented with *p-*value and 95% confidence interval (CI). The results reveal that U5MR for SC children belonging to Bihar, Jharkhand, Madhya Pradesh, Rajasthan and Uttar Pradesh are substantially higher than that of Non-SC/ST children. On the other hand, U5MR of ST children belonging to Chhattisgarh, Jharkhand, Madhya Pradesh, Orissa and Rajasthan are relatively higher than the corresponding U5MR of non-SC/ST population.

**Table 1.** U5MR (per 1000 live births) in five years preceding the survey for SC,ST and Non-SC/ST population in high focus states of India, 2015-16

| High focus states | SC |  |  |  | ST |  |  |  | Non-SC/ST |  |  |  |
| --- | --- | --- | --- | --- | --- | --- | --- | --- | --- | --- | --- | --- |
|  | U5MR | P value | 95% CI |  | U5MR | P value | 95% CI |  | U5MR | P value | 95% CI |  |
|  |  |  | Lower | Upper |  |  | Lower | Upper |  |  | Lower | Upper |
| Assam | 50.3 | 0.000 | 31.9 | 68.7 | 51.0 | 0.000 | 39.3 | 62.7 | 58.1 | 0.000 | 52.9 | 63.4 |
| Bihar | 72.9 | 0.000 | 65.0 | 80.9 | 52.5 | 0.000 | 38.7 | 66.2 | 54.0 | 0.000 | 50.2 | 57.8 |
| Chhattisgarh | 56.4 | 0.000 | 40.1 | 72.8 | 80.0 | 0.000 | 69.5 | 90.6 | 56.5 | 0.000 | 47.0 | 65.9 |
| Jharkhand | 59.5 | 0.000 | 47.7 | 71.2 | 63.8 | 0.000 | 55.2 | 72.4 | 48.6 | 0.000 | 42.0 | 55.2 |
| Madhya Pradesh | 68.5 | 0.000 | 62.4 | 74.6 | 78.3 | 0.000 | 71.5 | 85.0 | 57.8 | 0.000 | 53.4 | 62.2 |
| Orissa | 45.3 | 0.000 | 35.4 | 55.1 | 65.1 | 0.000 | 53.8 | 76.5 | 39.7 | 0.000 | 33.9 | 45.4 |
| Rajasthan | 61.8 | 0.000 | 54.0 | 69.5 | 57.8 | 0.000 | 50.2 | 65.5 | 45.3 | 0.000 | 41.1 | 49.4 |
| Uttar Pradesh | 85.5 | 0.000 | 79.9 | 91.1 | 58.5 | 0.000 | 37.0 | 80.1 | 76.0 | 0.000 | 72.5 | 79.4 |
| Uttarakhand | 43.8 | 0.000 | 32.9 | 54.7 | 35.9 | 0.004 | 11.8 | 60.0 | 48.0 | 0.000 | 41.3 | 54.7 |
| **India** | **55.8** | **0.000** | **53.7** | **58.0** | **57.2** | **0.000** | **53.7** | **60.8** | **46.6** | **0.000** | **45.4** | **47.9** |

### District level variation in U5MR by caste group in the districts of high focus states

We present district-level estimates of U5MR (S1 Table and Fig 2) for the SC, ST, non-SC/ST and total population separately. Each estimated value is supported by a *p* value and a 95% confidence interval to indicate the statistical significance of the U5M estimate. S2 Fig presents district level estimates of U5MR by caste groups. While 59% (105 of 179 districts with significant estimates) of the districts have higher mortality for the ST population (higher than 70 per 1000 live births), only 29% (88 of 302 districts with significant estimates) of districts have higher mortality for the non-SC/ST population. The corresponding figure for the SC population is 52% (148 of 283 districts with significant estimates). On the other hand, 14% (41 out of 283) of SCs, 18% (32 out of 179) of STs and 33% (101 out of 302) of non-SC/STs belong to districts with an U5MR range of below 50. Our findings show that districts performing poorly in terms of U5MR for the SC population are geographically concentrated in the states of Rajasthan, Uttar Pradesh, Bihar, and Madhya Pradesh, whereas for STs populations, they are concentrated in Madhya Pradesh, Chhattisgarh, and Orissa.

### Socioeconomic differentials of each caste group

Table 2 shows differences in the selected demographic, socioeconomic and program indicators from SCs, STs and the non-SC/ST population. U5M is higher among SCs and STs compared to the non-SC/ST group. While the mothers of approximately 40% of the SC and ST children have their first child before the age of 20 years, only 35% of the mothers of non-SC/ST children gave birth for the first time before age 20. There are more children with a birth order of four or higher among the SC (23%) and ST (20%) population compared to that for non-SC/ST children (18%). We also observed that the mother’s level of education varies widely by caste group. The number of parents with a secondary and higher level of educational attainment is lower for the parents of SC/ST children than for the parents of non-SC/ST children. About 71% of SC and 83% of ST children belong to poor households compared to only 53% of non-SC/ST children. Similarly, about 85% of SC and 93% of ST children live in rural areas, whereas 79% of the non-SC/ST children are rural inhabitants.

**Table 2.** Comparison of selected characteristics of children under five by social groups of high focus states in India, 2015–16

| Characteristics | SC |  | ST |  | Non SC/ST |  | Total |  |
| --- | --- | --- | --- | --- | --- | --- | --- | --- |
|  | N | % | N | % | N | % | N | % |
| **Under-five mortality** |  |  |  |  |  |  |  |  |
| No | 28,781 | 93.8 | 15,525 | 94.4 | 93,940 | 94.7 | 1,38,247 | 94.5 |
| Yes | 1,890 | 6.2 | 928 | 5.6 | 5,264 | 5.3 | 8,082 | 5.5 |
| **Demographic indicators** |  |  |  |  |  |  |  |  |
| **Sex of the child** |  |  |  |  |  |  |  |  |
| Female | 14,846 | 48.4 | 8,103 | 49.3 | 46,982 | 47.4 | 69,931 | 47.8 |
| Male | 15,826 | 51.6 | 8,350 | 50.7 | 52,222 | 52.6 | 76,398 | 52.2 |
| **Mother's age at first birth (yr)** |  |  |  |  |  |  |  |  |
| <20 | 12,336 | 40.2 | 6,688 | 40.6 | 35,067 | 35.3 | 54,091 | 37.0 |
| 20-24 | 15,348 | 50.0 | 7,839 | 47.6 | 51,093 | 51.5 | 74,280 | 50.8 |
| ≥ 25 | 2,988 | 9.7 | 1,926 | 11.7 | 13,045 | 13.1 | 17,958 | 12.3 |
| **Birth order** |  |  |  |  |  |  |  |  |
| First | 10,369 | 33.8 | 6,176 | 37.5 | 38,157 | 38.5 | 54,702 | 37.4 |
| 2-3 | 13,219 | 43.1 | 7,073 | 43.0 | 43,193 | 43.5 | 63,484 | 43.4 |
| 4+ | 7,084 | 23.1 | 3,204 | 19.5 | 17,855 | 18.0 | 28,143 | 19.2 |
| **Socioeconomic indicators** |  |  |  |  |  |  |  |  |
| **Mother's level of education** |  |  |  |  |  |  |  |  |
| Primary or below | 19,388 | 63.2 | 11,106 | 67.5 | 51,176 | 51.6 | 81,670 | 55.8 |
| Secondary | 9,669 | 31.5 | 4,833 | 29.4 | 38,328 | 38.6 | 52,830 | 36.1 |
| Higher | 1,615 | 5.3 | 514 | 3.1 | 9,701 | 9.8 | 11,829 | 8.1 |
| **Father's level of education<sup>s</sup>** |  |  |  |  |  |  |  |  |
| Primary or below | 2,246 | 40.4 | 1,598 | 49.3 | 6,167 | 33.3 | 10,011 | 36.6 |
| Secondary | 2,827 | 50.8 | 1,418 | 43.7 | 9,491 | 51.3 | 13,736 | 50.3 |
| Higher | 489 | 8.8 | 227 | 7.0 | 2,853 | 15.4 | 3,570 | 13.1 |
| **Type of residence** |  |  |  |  |  |  |  |  |
| Urban | 4,678 | 15.3 | 1,242 | 7.5 | 21,309 | 21.5 | 27,228 | 18.6 |
| Rural | 25,994 | 84.7 | 15,211 | 92.5 | 77,896 | 78.5 | 1,19,101 | 81.4 |
| **Wealth quintile (household)** |  |  |  |  |  |  |  |  |
| Poor | 21,623 | 70.5 | 13,579 | 82.5 | 52,108 | 52.5 | 87,310 | 59.7 |
| Middle | 4,382 | 14.3 | 1,508 | 9.2 | 16,810 | 16.9 | 22,701 | 15.5 |
| Rich | 4,666 | 15.2 | 1,366 | 8.3 | 30,286 | 30.5 | 36,318 | 24.8 |
| **Type of fuel used for cooking<sup>#</sup>** |  |  |  |  |  |  |  |  |
| Clean fuel & food not cooked | 4,249 | 14.8 | 1,052 | 6.7 | 24,069 | 26.1 | 29,371 | 21.5 |
| Solid fuel | 24,503 | 85.2 | 14,602 | 93.3 | 68,212 | 73.9 | 1,07,317 | 78.5 |
| **Type of toilet<sup>#</sup>** |  |  |  |  |  |  |  |  |
| Improved toilet | 7,363 | 25.6 | 2,665 | 17.0 | 40,651 | 44.1 | 50,680 | 37.1 |
| Not improved toilet | 21,389 | 74.4 | 12,989 | 83.0 | 51,630 | 55.9 | 86,007 | 62.9 |
| **Source of drinking water<sup>#</sup>** |  |  |  |  |  |  |  |  |
| Improved water | 26,891 | 93.5 | 12,529 | 80.0 | 86,185 | 93.4 | 1,25,605 | 91.9 |
| Not improved water | 1,862 | 6.5 | 3,125 | 20.0 | 6,096 | 6.6 | 11,083 | 8.1 |
| **Program indicators** |  |  |  |  |  |  |  |  |
| **Place of birth** |  |  |  |  |  |  |  |  |
| Home delivery | 9,132 | 29.9 | 5,747 | 35.0 | 25,535 | 25.8 | 40,414 | 27.7 |
| Institutional delivery | 21,432 | 70.1 | 10,657 | 65.0 | 73,380 | 74.2 | 1,05,468 | 72.3 |
| **Antenatal visits during pregnancy (last birth)** |  |  |  |  |  |  |  |  |
| None | 5,894 | 27.6 | 2,914 | 24.5 | 16,164 | 22.6 | 24,972 | 23.8 |
| 1 visit | 1,821 | 8.5 | 682 | 5.7 | 5,437 | 7.6 | 7,940 | 7.6 |
| 2-3 visits | 7,972 | 37.3 | 4,106 | 34.5 | 25,744 | 36.0 | 37,822 | 36.1 |
| 4+ visits | 5,672 | 26.6 | 4,197 | 35.3 | 24,241 | 33.9 | 34,110 | 32.5 |
| **Total** | **30,671** | **100** | **16,453** | **100** | **99,204** | **100** | **1,46,329** | **100** |

<sup>§</sup>Excluded missing cases. <sup>#</sup>Not included de jure resident. All percentages are weighted cases
All the covariates are tested with Pearson chi-squared test and found to be statistically significant at $p < 0.001$

In terms of environmental factors, the use of solid fuel for cooking is higher among the households of SC (85%) and ST (93%) children compared to those of non-SC/ST (74%) children. The use of unimproved sanitation facilities and unsafe sources of drinking water are substantially higher among ST children compared to non-SC/ST children. As far as program indicators are concerned, the proportion of institutional delivery is lower and the number of “never visit” responses for ANC during pregnancy is higher among SCs and STs compared to those of the non-SC/ST population in the high focus states of India.

### Association between U5M and different demographic, socioeconomic and program indicators

The association between U5M and different demographic, socioeconomic and program indicators are presented in the logistic regression model in Table 3. In model 1, we present the unadjusted effect of caste on U5M. The results show that compared to non-SC/ST children, SC (OR: 1.19,p<0.01, SE:0.03) and ST (OR: 1.10,p<0.05,SE:0.03) children have a higher likelihood of dying. After controlling for other socioeconomic factors in model 2, children belonging to SCs have a 1.15 times higher likelihood of dying. We did not find a statistically significant coefficient for children belonging to scheduled castes in model 2.

**Table 3.** Logit estimates of U5M by different characteristics of high focus states in India, 2015-16

| Variables | U5M |  |
| --- | --- | --- |
|  | Unadjusted OR<br>(Model 1) | Adjusted OR<br>(Model 2) |
| **Demographic indicators** |  |  |
| **Sex of the child** |  |  |
| Female ® |  |  |
| Male |  | 1.01(0.03) |
| **Mother's age at first birth (yr)** |  |  |
| <20® |  |  |
| 20-24 |  | 0.97(0.03) |
| ≥ 25 |  | 1.13**(0.06) |
| **Birth order** |  |  |
| 1 ® |  |  |
| 2-3 |  | 0.76*** (0.03) |
| 4+ |  | 1.19*** (0.05) |
| **Socioeconomic indicators** |  |  |
| **Caste** |  |  |
| Non-SC/ST® |  |  |
| SC | 1.19*** (0.03) | 1.15*** (0.05) |
| ST | 1.10** (0.03) | 0.93(0.04) |
| **Mother's level of education** |  |  |
| Primary or below ® |  |  |
| Secondary |  | 0.83*** (0.03) |
| Higher |  | 0.68*** (0.06) |
| **Father's level of education** |  |  |
| Primary or below ® |  |  |
| Secondary |  | 0.75*** (0.06) |
| Higher |  | 0.68*** (0.10) |
| **Type of residence** |  |  |
| Urban® |  |  |
| Rural |  | 1.02(0.05) |
| **Wealth quintile (household)** |  |  |
| Poor ® |  |  |
| Middle |  | 0.90** (0.05) |
| Rich |  | 0.75*** (0.04) |
| **Type of fuel for cooking** |  |  |
| Clean fuel & food not cooked ® |  |  |
| Solid fuel |  | 0.93(0.05) |
| **Type of toilet** |  |  |
| Improved toilet ® |  |  |
| Not improved toilet |  | 1.14*** (0.05) |
| **Source of drinking water** |  |  |
| Improved water ® |  |  |
| Not improved water |  | 0.93(0.05) |
| **Program indicators** |  |  |
| **Place of delivery** |  |  |
| Home delivery® |  |  |
| Institutional delivery |  | 0.92** (0.03) |
| **ANC visits during pregnancy (last birth)** |  |  |
| None ® |  |  |
| 1 visit |  | 0.87** (0.06) |
| 2-3 visits |  | 0.85*** (0.04) |
| 4+ visits |  | 0.73*** (0.03) |
| **Constant** | 0.05*** (0.01) | 0.07*** (0.01) |
| **Observation** | **1,46,329** | **113,325** |

Notes: OR-Odds ratio; (a) \*\*\* $p < 0.01$ , \*\* $p < 0.05$ , \* $p < 0.1$ (b) The entries in parenthesis refer to standard errors (c) ® indicate reference category

The regression results also revealed that parents’ level of educational attainment has a statistically significant effect on reducing U5M and as expected, U5M shows significant declines with increases in the educational level of the parents. Children are less likely to die when their parents have completed secondary and higher education than children born to parents who have a primary level of education or lower. Similar results were observed for household wealth as for educational attainment. As the level of wealth of the household increases, the odds of U5M significantly decreases. It is important to note that the effect of parental education in the reduction of U5M is more rigorous than the effect of the household wealth quintile. The odds ratio for children of a mother with a secondary level education is 0.83 (p<0.01, SE:0.03) versus the odds ratio for children belonging to the middle wealth quintile, which is 0.90 (p<0.05, SE:0.05). The odds ratio for children of highly educated mothers is 0.68 (p<0.01, SE:0.06) versus an odds ratio for children that belong to the “rich” quintile, which is 0.75(p<0.01, SE:0.04).

The age of mothers at the birth of their first child was found to be significantly associated with U5M. Children born to women aged 25 years or older had significantly higher odds (OR: 1.13, p<0.05, SE:0.06) of child mortality than babies born to women younger than 20 years. Birth order was significantly associated with U5M and the results show that children with a birth order of four or more (OR: 1.19, p<0.01, SE:0.05) have a higher likelihood of dying before the age of five. Male children had slightly higher odds of dying before their fifth birthday compared to female children, but the finding was not statistically significant.

Among the environmental factors, the type of toilet used is significantly associated with U5M. The likelihood of U5M is higher among children from households with non-improved toilet facilities (OR: 1.14, p<0.01, SE:0.05) compared to children of households with improved toilet facilities. Place of birth and number of ANC visits during pregnancy were found to be significant indicators in reducing U5M in the high focused states of India. The likelihood of lower U5M was also increased when mothers were able to opt for institutional delivery (OR: 0.92,p<0.05, SE:0.03) rather than home delivery. The findings also show that as the frequency of ANC visits increase, the odds of U5M decreases significantly.

### Results of the decomposition analysis

The results of the detailed decomposition are presented in Table 4. To make our results more readable, we present the coefficient in terms of percentages in Figure 3. The positive contribution of a covariate indicates that this particular covariate contributed to widening the U5M gap between SCs/STs and the non-SC/ST populations, while the negative contribution of a covariate indicates that it was helping to reduce the gap. Our results indicate that 73% of the U5M gap between SCs/STs and the non-SC/ST population can be explained by the factors included in the analysis. The unexplained gap (27%) for U5M might be related to other structural factors not covered by the dataset in the analysis.

**Table 4.** Results of Fairlie decomposition of average gap in U5M risk between caste groups in the high focus states of India, 2015-16

| Covariates | U5M |
| --- | --- |
|  | Contribution |
| Mother's level of education | 0.934*** |
| Father's level of education | 0.001 |
| Mother's age at first birth | -0.143 |
| Sex of the child | -0.140** |
| Birth order | -0.069 |
| Wealth of the household | 2.885*** |
| Type of fuel used for cooking | 0.403 |
| Type of toilet | -0.081 |
| Source of drinking water | -0.270** |
| Type of residence | 0.200 |
| Place of delivery | 0.566*** |
| ANC visits | 0.610*** |
| **Total gap** | **6.69** |
| **Explained gap** | **4.90 (73%)** |
| **Number of observations** | **1,13,325** |

Note: (a)\*\*\* p< 0.01, \*\* p< 0.05 & \* p< 0.10

**Fig 3.**
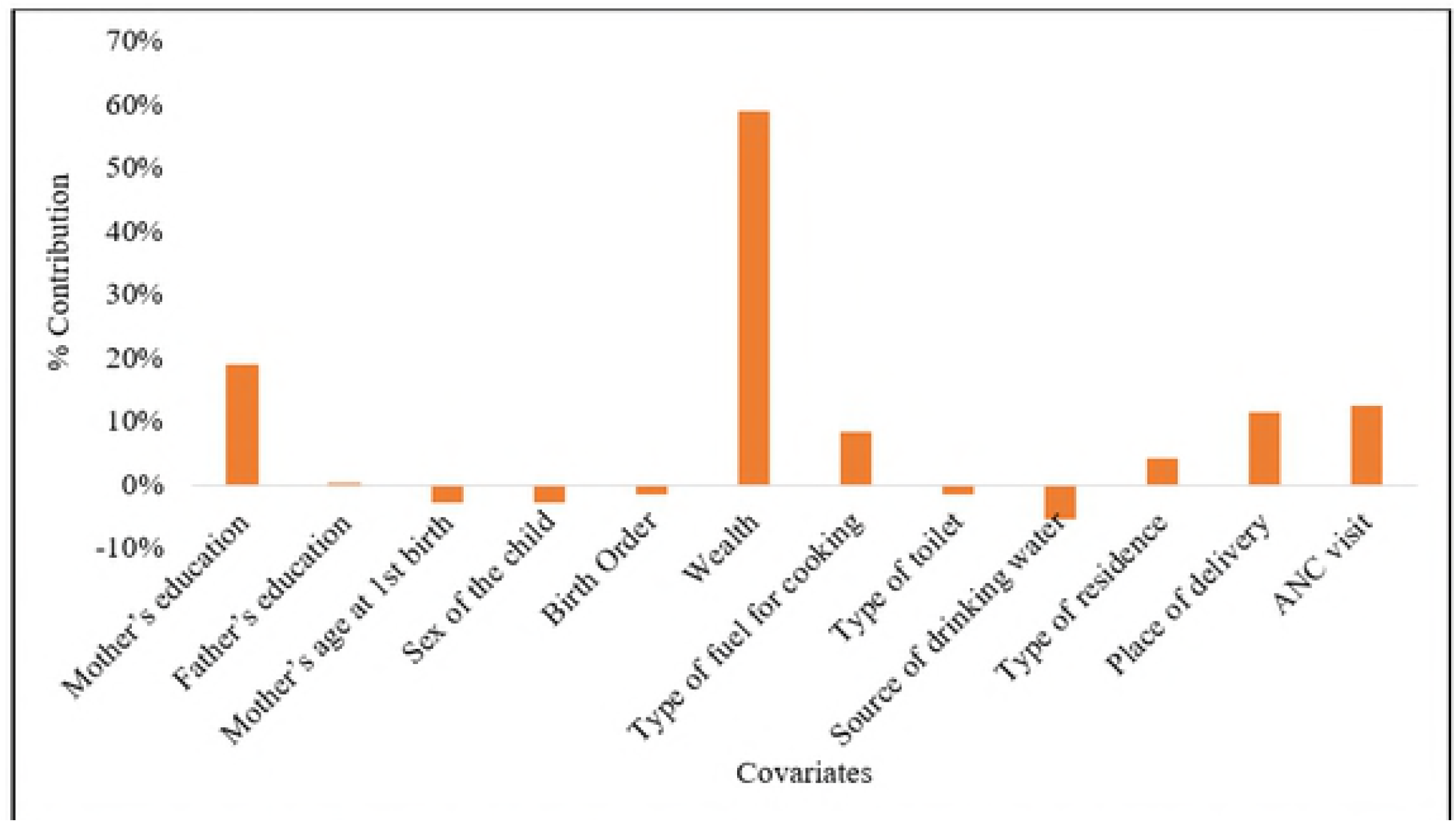
Results of Fairlie decomposition analysis showing percentage contribution of each covariate to the gap in U5M-risk between caste groups in high focused states of India, 2015–16

Of the explained gap, 78% can be related to differences in the distribution of women’s educational attainment and household wealth for U5M risk. Household wealth is the most significant contributor (59%) to the gap in U5M between SCs and STs followed by the mother’s level of educational attainment (19%). Program indicators such as place of delivery and number of ANC visits also contribute (12% each respectively) to widening the U5M gap between SCs/STs and the non-SC/ST population at a 10% level of significance. Demographic variables such as the sex of the child, the mother’s age at the birth of her first child and birth order contributed to reducing the gap. While the source of drinking water narrowed the gap (9%), the type of toilet used in the household diminished the caste gap in U5M, although not significantly. The type of fuel used for cooking and the type of residence widened the caste gap in U5M risk, although its contributions are not statistically significant.

## Discussion and conclusion

Any country’s general medical and public health conditions, and consequently its level of socioeconomic development, can be measured based on the health of its children. Although U5M has been continuously declining in India in recent decades, it is still substantially higher among certain social groups and regions in the country, particularly before the implementation of the NRHM. This study documented disparities in U5M by caste groups in the high mortality regions of India in recent years, using nationally representative data. The novelty of the study lies therein that, to our knowledge, this is the first study in India that provides district level estimates of U5M and systematically investigates the factors explaining U5MR by caste groups using the most recent DHS data. This is also the first study to document the association between caste and U5M in the post-NRHM period in India.

Our study highlights a few important findings. First, the disparity in U5M, was profound by caste groups during the pre-NRHM period, and has been reduced drastically in recent years. This success may be attributed to the NRHM under which special provision has been made for maternal and child health care services for women and children belonging to deprived castes. For instance, under the NHM, a conditional cash transfer scheme was introduced for institutional deliveries. In the high performing states of India, this cash was given only to women from deprived caste groups, whereas in low performing states, this cash was available to all poor as well as deprived castes women [39–41]. Despite the reduction in the parallel gap in U5M between caste groups over time, our study further reveals that the caste gap in U5MR is persistent even in the NRHM/NHM period. Children belonging to deprived castes have a higher likelihood of dying than those belonging to non-deprived castes. Our analysis shows that this association remains significant for children from SCs even after controlling for other background characteristics. Thus, our study reconfirms that children belonging to deprived castes are still in a disadvantaged position in terms of mortality outcomes. These findings are consistent with those of previous national and sub-national studies [16,17,21,42]. Our results also indicate that macro level indicators of U5M at district level shows that the U5MR of the SC/ST population is higher than that of the non-SC/ST population. We observed a geographical clustering of U5MR in the studied area.

Secondly, this study contributes to the debate on whether maternal education or household economic status is more conducive for child mortality reeducation. There are numerous previous studies that documents the negative and strong role of education on mortality through various mechanisms such as increased knowledge, better accessibility, use of modern health care facilities, higher mobility etc.[43–47]. Yet, there is continuous debate around whether education should be prioritized over wealth for preventing under-five deaths. While many previous studies have described the independent effect of both maternal education and wealth status of the household on child mortality, some studies established that maternal education is the single most important determinant of child survival at all levels and its effects on child survival are stronger than those of household wealth [48–50]. A study based on 42 developing countries, however, argued that although higher education levels were associated with disproportionately greater returns to child health, the pattern for household wealth was erratic: in many countries there were diminishing returns to child health at higher levels of household wealth [51]. Our study in India’s high mortality areas support previous findings that maternal education is more important in the reduction of U5M than household wealth status.

The decomposition analysis allows us to gain insight into the relative contribution of various factors in the caste-based gaps in the U5MR. It demonstrates that the current gap in U5M by caste groups is mainly due to their disadvantages in terms of the economic conditions and educational status of their parents. Unlike the results from the regression analysis, household economic status contributes the most to the gap, followed by parental education. This indicates that there is a greater economic divide between well-off and deprived caste groups, which explains the majority of the gap in U5M between caste groups. Since more than 50% of SC/ST households are from the poorest backgrounds, it is not surprising that household economic status turned out to be the largest contributing factor in widening the caste gap in terms of U5M. It is argued that poor SC/ST households do not have enough resources for child and maternal health care expenses. In contrast, the non-SC/ST population is economically better off and more educated. They may have a more advanced view, more knowledge about child care and preventive care (greatly associated with the modern healthcare system), as well as higher confidence in dealing with health care providers and a greater ability and readiness to travel outside the community for their health needs [52], all of which may help to reduce poor child health outcomes.

Following the wealth status of the household, maternal educational attainment contributes significantly to widening the caste-based gap in U5M. This is consistent with the findings of previous research [23,53–55]. It is possible that the lower level of education among SCs and STs is accompanied by a low awareness of health services. This includes less knowledge of the benefits of preventive child health care, use of traditional health care, poor communication with the husband and other family members on health-related issues and poor decision-making power within the family, low self-confidence, poor survival abilities and poor negotiating skills with health care providers [56]. The type of cooking fuel used by households also positively contributed to widening the gap, which is an indication of less access to clean fuel by the disadvantaged caste groups. It is interesting to note that the two program related variables together contributed 24% to the gap in the U5MR. This is an interesting finding from a policy perspective as it shows that there is still scope to uplift access to health facilities for women and children belonging to deprived caste groups in the study area.

Despite the U5MR of our study area experiencing a faster decline in the most recent years compared to the stagnation in mortality reduction that was observed in the early 2000s [57], this reduction is still not enough to achieve the SDG goals on preventable neonatal and under-five deaths [3]. Previous findings indicate that nearly all districts of Uttar Pradesh, Bihar, Madhya Pradesh, and Chhattisgarh will fail to achieve the SDG3 goal on neonatal mortality rate [3]. Similarly, in Uttar Pradesh, not a single district is expected to meet the target for U5MR as set out in SDG3 [3]. Since our findings demonstrate that maternal education is relatively more effective in U5MR reduction, there is a greater need to raise the level of educational attainment, particularly secondary education among girls in the high focus states of India. At the same time, programs uplifting the economic status of disadvantaged groups is equally important for a faster and sustainable reduction of U5MR in the high focus states. To improve universal coverage and access to maternal and child health care services, emphasis should be placed on creating awareness of the district level intervention program through community-based awareness programs, as well as on educating parents about the possible high-risk factors and preventive measures associated with child health [3]. Another possible initiative might be to involve the parents of SC and ST children in health-related intervention programs at village or community level and educate them about preventive care for their infants at home and the importance of antenatal care for women during pregnancy. In addition, sensitization to and creating awareness around preventive health care, maternal care, nutrition, awareness about infectious diseases, the benefits of hygiene and sanitation, and subsidized maternal health care services among SC and ST populations, should be increased through outreach programs.

## Supporting information

**S1 Table. Estimated Districtwise Under-five mortality rate for ten-years periods preceding the survey for SC, ST, Non-SC/ST and Total in high focus states of India, 2015-16. (PDF)**

**Appendix S1. Description of Fairlie method. (DOC)**

